# RNA accessibility sequencing reveals transcriptome-wide RBP binding and RNA structure

**DOI:** 10.64898/2026.11.02.763788

**Authors:** Yang Li, Chang Ye, Ruiqi Ge, Cheng-Wei Ju, Yu Xiao, Weixin Tang, Mengjie Chen, Chuan He

## Abstract

We introduce RAD-seq (RNA Accessibility via Deamination sequencing), a simple and scalable method that quantifies RNA accessibility by protein binding in vivo. RAD-seq uses TadA8.20-mediated adenosine deamination to report the local exposure of each adenosine to its molecular environment, integrating the combined effects of base pairing, RNA folding, and RNA-binding protein (RBP) occupancy in a single experiment. RAD-seq profiles recapitulate known structural signatures, reveal high-resolution RBP footprints, report on the influence of RNA modifications, and capture dynamic remodeling of RNA accessibility during oxidative and heat-shock stress. By integrating RAD-seq with RNA Bind-n-Seq motifs, we detected local enrichment of the RNA *N*^6^-methyladenosine (m^6^A) methyltransferase recruiter TRA2A following knockdown of the exon junction anchor protein EIF4A3, revealing a potential mechanism that links RNA accessibility to enhanced TRA2A recruitment and m^6^A deposition. Extending this framework, we introduce an unsupervised statistical model that combines RAD-seq with motif priors to infer binding landscapes for multiple RBPs, reproducing the information from numerous CLIP-seq datasets from a single RAD-seq experiment. These results indicate RAD-seq as a generalizable approach for joint profiling of RNA structure and protein occupancy, defining enzymatic RNA accessibility as a new dimension to investigate regulatory elements on RNA.

## Introduction

The concept of chromatin accessibility transformed our understanding of gene regulation by providing a quantitative measure of DNA’s openness to regulatory factors. Assays such as DNase-seq^1^ and ATAC-seq^2^ developed this idea into a unified readout that captures enhancer activity, transcriptional potential, and cell identity. The ability to simultaneously capture the effects of these disparate influences offered an unprecedented ability to observe the dynamics of chromatin organization and the regulatory factors shaping it. Using this new measure, researchers could rapidly nominate genomic regions with high regulatory potential, quantify the effects of specific regulatory factors on genome-wide transcriptional openness, or identify genetic variants whose effects were mediated through shifts in chromatin accessibility. However, no comparable, nucleotide-level metric has been established for RNA, despite its central role in gene expression control. Associations between RNA molecules and other components of the genome, transcriptome, and cellular machinery serve critical functions in virtually every step along the pathway from genetic code to protein expression. Yet targeted assays are currently used to test specific aspects of its structural conformation or RBP binding instead of a more integrated measure of *RNA accessibility*, which would allow for discovery-oriented profiling of potential RNA regulatory elements in the whole transcriptome scale.

In contrast to chromatin accessibility, which can be straightforwardly defined by steric openness to trans-acting proteins, RNA accessibility encompasses multiple influences, including base pairing, RNA-binding proteins (RBPs), and RNA folding. Existing methods provide fragmented views of various components of RNA accessibility. Chemical probing approaches such as Keth-seq^3^, DMS-seq^4^, Structure-seq^5^, SHAPE-seq^6^, icSHAPE^7^, and corresponding mutational profiling approaches^8–10^ have been widely used to map base-pairing by exploiting the selective reactivity of single-stranded nucleotides. Other chemical methods like LASER-seq, LASER-MaP^11,12^ and icLASER^12^ probe RNA solvent accessibility in cells. These methods elegantly reveal secondary structure or solvent accessibility landscapes, but because they rely on small probes, they may be less sensitive to protein-scale steric occlusion, conceptually analogous to the difference between fine-scale chemical footprinting and accessibility-based chromatin profiling methods. In addition, high sequencing depth is typically required to achieve adequate coverage across transcripts because of the limitation of reverse-transcription-stop-based methods or the limitation of error rate required for reliable mapping in mutational profiling (MaP) approaches. Conversely, nuclease-based assays like PARS^13,14^, FragSeq^15^, and PIP-seq^16,17^ detect protection patterns that combine structural and protein effects but do so at low resolution and with limited robustness. Meanwhile, protein-centric assays, including CLIP-seq^18,19^, iCLIP^20^, and eCLIP^21^, focus on individual RBPs and rely on immunoprecipitation or tagging—requiring one experiment per protein and offering incomplete coverage. More recent global strategies, such as RNP-MaP^22^ can profile protein–RNA interactions transcriptome-wide by using a hetero-bifunctional crosslinker to freeze interacting proteins in place on RNA and read the signal by MaP. Collectively, these approaches have generated a wealth of insights but have not converged on a conceptual or operational definition of RNA accessibility that integrates structure and RBP occupancy into a single measurable property.

To fill this gap, we introduce RAD-seq (RNA Accessibility via Deamination sequencing), an assay that quantifies enzymatic RNA accessibility or RNA protection by secondary structure and protein binding. While RNA deaminases have been fused with different RBPs to map their binding sites^23–28^, RAD-seq leverages the evolved tRNA adenosine deaminase TadA8.20^29–31^ as an enzymatic probe of RNA accessibility. TadA8.20 deaminates exposed adenines (A), converting them to inosines (I) that are read as guanosines (G) in sequencing. The rate of A→I conversion thus encodes the local accessibility of each adenine, integrating both secondary structure and RBP occlusion^29^. Because TadA catalysis requires physical access to the RNA bases, reduced conversion marks either base-paired or protein-bound nucleotides. In this way, RAD-seq produces a direct single-nucleotide resolution and transcriptome-wide measure of RNA enzymatic accessibility.

A key innovation of RAD-seq is its dual-mode design. By comparing “native” samples, where TadA senses structural openness in vivo, with “fixed” samples, where mild formaldehyde fixation preserves RNA–protein contacts, RAD-seq distinguishes the contributions of structure and protein occupancy. The difference between native and fixed accessibility effectively isolates the RBP-occluded fraction, transforming RAD-seq into a dual readout of RNA structure and protein binding within the same experimental framework. This approach parallels ATAC-seq’s measurement of chromatin openness to transposases, as well as DNA accessibility measurements through 5mC and 6mA methylation introduced by DNA methyltransferases^32–36^, extending the core principle into the RNA domain with molecular specificity. The simplicity and efficiency of this method, which returns accurate transcriptome-wide accessibility profiles based on read counts an order of magnitude lower than many comparable methods, make it a highly cost-effective and scalable alternative to existing methods of probing RNA’s openness to biochemical interactions.

In addition to its experimental innovation in quantifying raw RNA accessibility, RAD-seq data can be integrated with computational modeling to infer RBP occupancy transcriptome-wide. Accessibility features, combined with motif enrichment, conservation, and expression, accurately reconstruct RNA–protein interaction landscapes. We show that RAD-seq footprints align with eCLIP peaks, capture known binding motifs, and reveal condition-dependent occupancy changes. A motif-guided probabilistic model further infers transcriptome-wide binding probabilities for dozens of RBPs simultaneously, compressing what would otherwise require hundreds of CLIP-seq experiments into a single assay.

Together, these advances define enzymatic RNA accessibility as a measurable molecular property and establish RAD-seq as its defining technology. Like ATAC-seq for chromatin, RAD-seq unifies RNA structure and protein occupancy into a single, quantitative readout, enabling a global and mechanistically interpretable view of RNA-mediated regulation. Beyond its immediate applications to structure probing and RBP mapping, RAD-seq offers a general framework for linking RNA conformation, protein occupancy, and gene regulation, while remaining read-efficient, scalable, and adaptable to diverse biological contexts even at modest sequencing depth.

## Results

### RNA accessibility and the principle of RAD-seq

Chromatin accessibility quantifies DNA openness to nuclear factors, establishing a conceptual framework for how physical accessibility encodes regulatory potential. In parallel, RNA accessibility reflects the exposure of individual nucleotides to molecular interactions, jointly determined by base pairing and RNA-binding protein (RBP) occupancy (Fig. 1a). To capture this property in vivo, we employed the evolved adenosine deaminase TadA8.20 as an enzymatic reporter of local accessibility. In RAD-seq (RNA Accessibility via Deamination sequencing), TadA8.20 converts accessible adenines to inosines, which are read as guanosines during sequencing, yielding a single-nucleotide accessibility score (AS) defined as the fraction of G calls among reads covering each adenosine (Fig. 1b).

**Figure 1.**
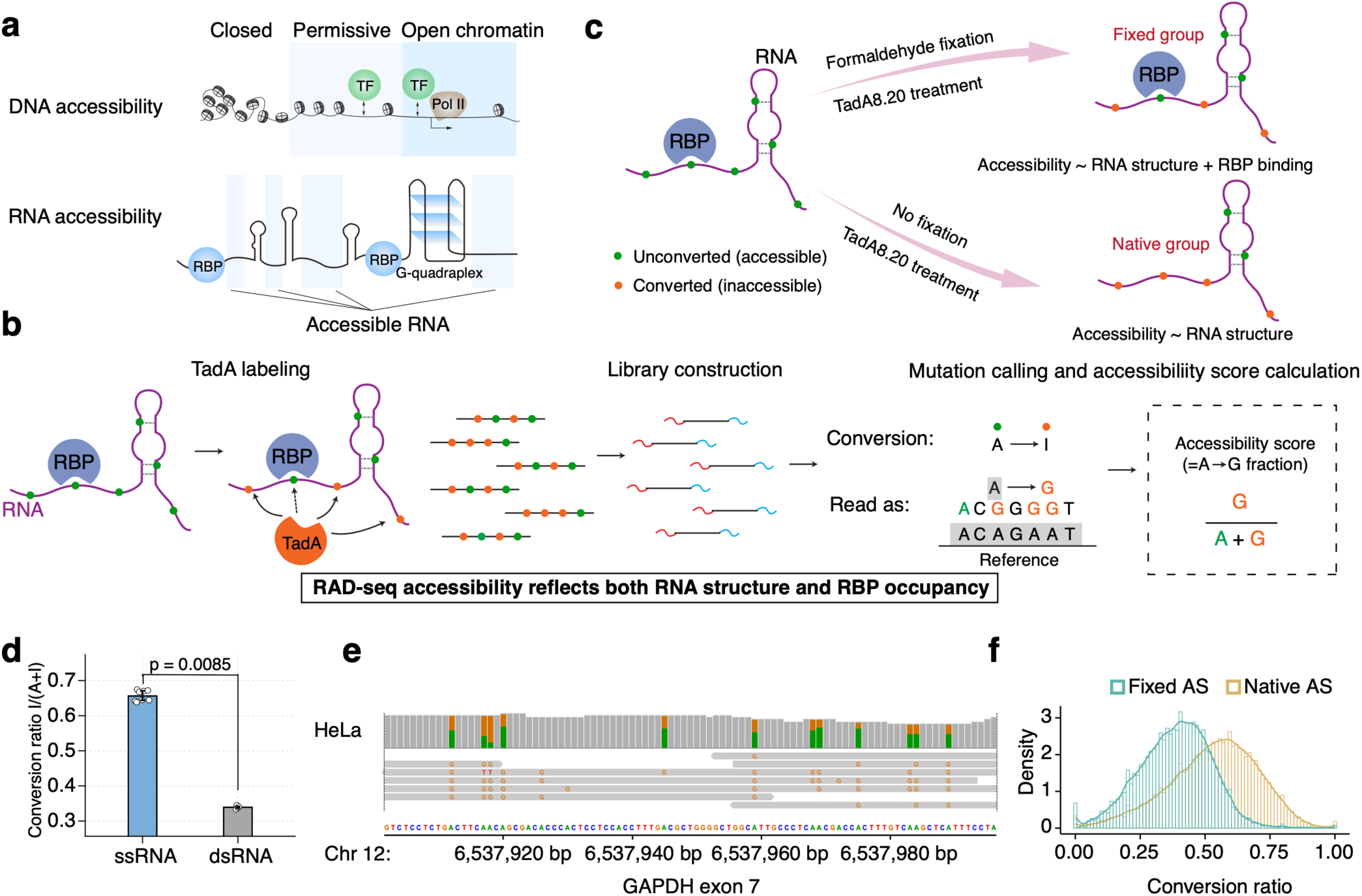
Overview and validation of RAD-seq for transcriptome-wide RNA accessibility. (a) Conceptual overview of DNA (chromatin) accessibility versus RNA accessibility. DNA accessibility is shaped by chromatin structure and transcription factor binding; RNA accessibility reflects RNA structure and RNA-binding protein (RBP) occupancy. The DNA accessibility schematic is adapted from Klemm *et al*.^56^ (b) RAD-seq workflow. TadA8.20 catalyzes in vivo A→I deamination at unprotected adenosines. After library preparation and sequencing, inosines are read as G, and the A→G conversion ratio defines the accessibility score (AS). (c) Formaldehyde crosslinking distinguishes native AS (RNA structure only) from fixed AS (RNA structure plus RBP occupancy). (d) Validation on synthetic RNA oligonucleotides by LC-MS/MS (QQQ). Conversion ratios for ssRNA (single 77 nt oligo) and dsRNA (annealed complementary 77 nt oligos) are shown as medians with interquartile ranges; dots are individual replicates (n = 4 per group). Groups were compared with a two-sided Wilcoxon rank-sum test (p = 0.0085). (e) IGV tracks of A→G conversions across GAPDH exon 7. Robust AS estimates are obtained at relatively low sequencing depth due to high deamination efficiency. (f) Transcriptome-wide distributions of native and fixed AS (conversion ratios).

To distinguish structural openness from protein occlusion, we performed both native and formaldehyde-fixed reactions: fixation preserves RBP–RNA contacts prior to lysis so that the difference between conditions reveals the protein-bound fractions (Fig. 1c). Therefore, RAD-seq unifies measurements of RNA structure and protein occupancy in a single assay.

We first validated TadA8.20 reactivity using synthetic RNA oligonucleotides designed to adopt single-stranded or duplex conformations. The enzyme displayed markedly higher A→I conversion on single-stranded substrates than on duplexes (P = 0.0085; Fig. 1d), confirming sensitivity to base-pairing. Applying RAD-seq to HeLa cells, we observed robust A→G conversion across broad transcript regions under both native and fixed conditions, providing reproducible accessibility estimates at moderate sequencing depth (Fig. 1e,f). Accessibility profiles were highly concordant across technical and biological replicates for both rRNA and mRNA (Supplementary Fig. 1), demonstrating that RAD-seq captures intrinsic transcript features rather than experimental noise. Comparison of local motif preferences across RNA structure and accessibility profiling methods at adenosine-centered sites showed that RAD-seq, like other methods, can be influenced by flanking sequence context (Supplementary Fig. 2). These findings suggest that local sequence context contributes to measured accessibility in a method-dependent manner.

Collectively, these results show that RAD-seq profiles RNA accessibility in vivo, resolving structural organization and RBP protection within a single experiment. The assay delivers a unified, nucleotide-resolution view of RNA states and establishes a formal, scalable definition of enzymatic RNA accessibility.

### Validation of RAD-seq accessibility against chemical probing of cellular RNA structure

To validate that RAD-seq accessibility profiling reveals RNA structure information correctly, we benchmarked it against existing structure-probing methods in the HeLa cell line. We selected icSHAPE, which captures the same class of single-stranded reactivities measured by DMS- or SHAPE-related methods, as the representative chemical-probing benchmark because it is the most widely used transcriptome-scale method for mapping RNA structure *in vivo*. Native RAD-seq consistently demonstrated comparable performance to icSHAPE in detecting RNA secondary structural motifs. For example, comparing the profiles for SNORA73A and SNORD3A (two small nucleolar RNAs with previously predicted secondary structures by R2DT^37^), we found that both native RAD-seq accessibility scores and icSHAPE reactivity scores clearly distinguished paired from unpaired regions (Fig. 2a,b, Supplementary Fig. 3a,b). The divergence in scores between single- and double-stranded regions was substantially greater for SNORA73A than SNORD3A, perhaps reflecting differences in RBP binding or tertiary structure between these molecules, but this observation was consistent between RAD-seq and icSHAPE (Fig. 2b, Supplementary Fig. 3b).

**Figure 2.**
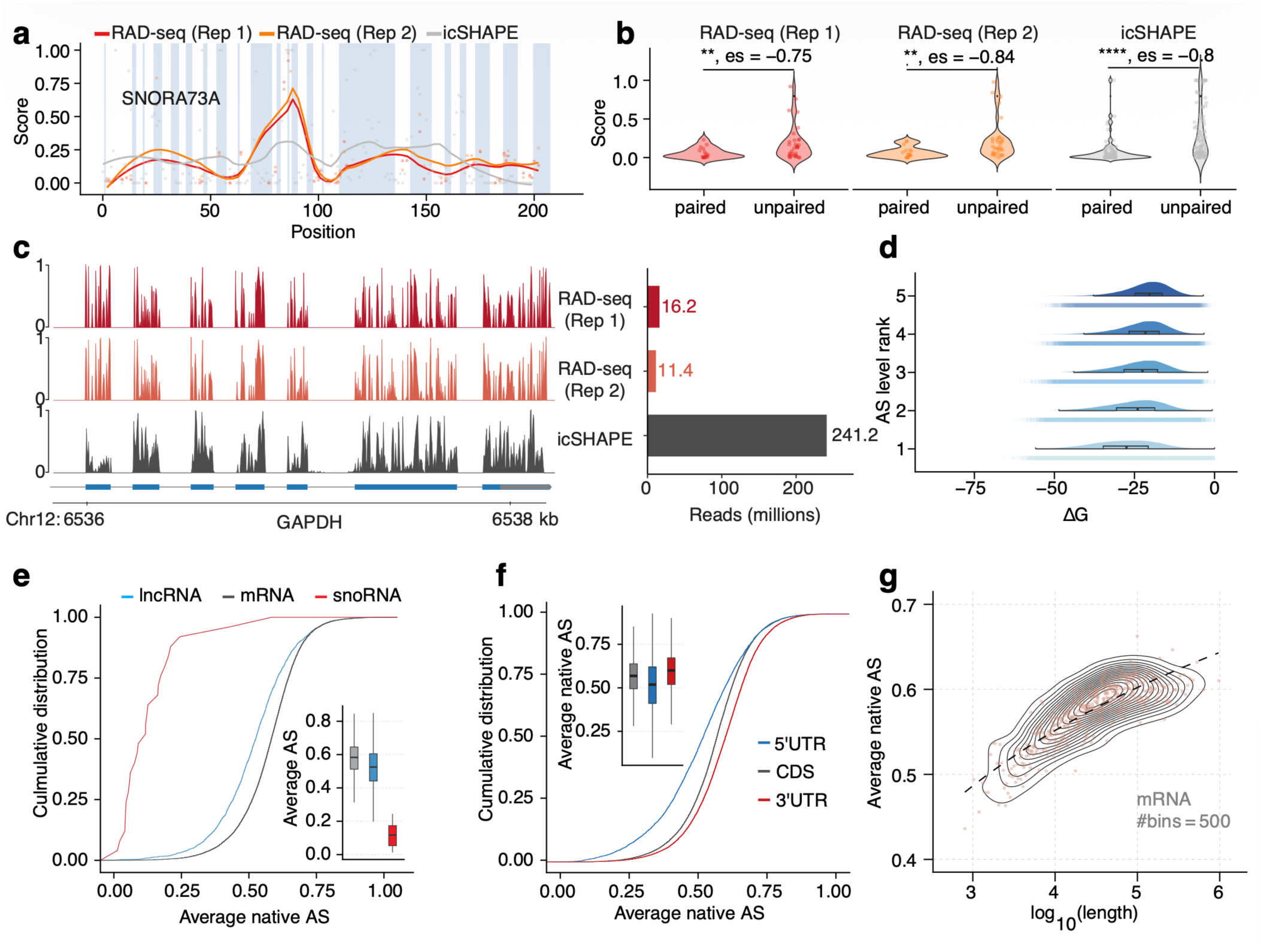
Transcriptome-wide RNA accessibility profile using RAD-seq. (a) Accessibility profile of SNORA73A shown in blue (n = 2 biological replicates) with the icSHAPE score in gray. Blue bands represent predicted double-stranded regions, while white bands represent predicted single-stranded regions. (b) Distributions of accessibility scores and icSHAPE scores across base-paired and unpaired regions of SNORA73A. (c) Accessibility profiles along the GAPDH transcript measured by RAD-seq (∼16 million reads) and icSHAPE (∼241 million reads), showing concordant long-range patterns. (d) Predicted RNA folding free energy (ΔG) across accessibility-score rank groups; values are binned into five equal-sized groups, where higher ΔG indicates a tendency toward more structured RNA as defined in Methods. The predicted RNA folding free energy data is downloaded from RNAStructuromeDB^38^. (e) Mean accessibility across protein coding, lncRNA, and snoRNA. (f) Cumulative distributions and boxplots of RNA accessibility in 5′ UTR, CDS, and 3′ UTR across the transcriptome. (g) Relationship between mean transcript accessibility and transcript length; Pearson r and two-sided P value from a linear fit are reported. Asterisks indicate significance as follows: ns, P ≥ 0.05; *, P < 0.05; **, P < 0.01; ***, P < 0.001; ****, P < 0.0001.

In addition to small RNAs, we compared short-range accessibility patterns in larger transcripts between native RAD-seq and icSHAPE and consistently observed close alignment. For example, both methods returned highly similar profiles for the exon 7 region of GAPDH despite RAD-seq’s nearly 20-fold lower read count (Fig. 2c). We also validated native RAD-seq accessibility scores across the whole transcriptome by comparing them to predictions of folding free energy (ΔG) from RNAstructuromeDB^38^ (Fig. 2d). Regions predicted to form stable structures (more negative ΔG) exhibited lower accessibility and higher Gini indices (Supplementary Fig. 4a,b), indicating that TadA8.20 deamination faithfully reports solvent exposure constrained by secondary structure. Collectively, our results demonstrate that native RAD-seq signal is consistent with predicted RNA secondary structure both at short range and on the level of the whole transcriptome, and that it aligns nucleotide-level measures of enzyme accessibility with established structure-probing assays. We therefore conclude that RAD-seq effectively captures a real property of RNA in living cells.

### RAD-seq accessibility reflects structural and functional characteristics of cellular RNA

Although RAD-seq is consistent with icSHAPE when profiling short-range features of the RNA structurome, we found that both native and fixed RAD-seq scores diverged from icSHAPE at the transcriptome level when assessed by per-site correlation (Supplementary Fig. 3c). Notably, both native and fixed RAD-seq scores demonstrated high correlation with icSHAPE when binned the sites into 500 bins (Supplementary Fig. 3d). These observations suggest that although RAD-seq accessibility signatures encompass the structural features detected by icSHAPE globally, they also capture additional aspects of RNA interaction locally, consistent with RAD-seq’s broader sensitivity to both base pairing and RBP protection rather than structure alone. The global divergence and local concordance between the two assays highlight RAD-seq’s unique ability to link RNA structure and occupancy within a single quantitative framework, complementing existing RNA structural probing methods such as icSHAPE.

Accessibility also varied by RNA biotype. Small nucleolar RNAs exhibited the lowest native accessibility, consistent with compact and protein-rich RNP assemblies (Supplementary Fig. 4c). Long noncoding RNAs (lncRNAs) displayed reduced native accessibility relative to mRNAs (Fig. 2e, Supplementary Fig. 4d), which likely implies more extensive structural stability and protein engagement. RAD-seq profile across the transcriptome revealed consistent positional patterns in both native and fixed accessibility scores, which peaked in 3′ UTRs and were lowest in 5′ UTRs whether quantified by mean accessibility or the Gini index (Fig. 2f, Supplementary Fig. 4e-g). These trends might indicate that areas near transcript 5’ ends are subject to greater structural constraints and experience more frequent regulatory protein interactions.

Transcript length further influenced accessibility in a biotype-specific manner. For mRNAs, longer transcripts showed significantly higher accessibility in both native and fixed conditions (P = 1.9 × 10⁻¹¹⁶; Fig. 2g, Supplementary Fig. 4h), which might imply reduced base pairing and decreased RBP association across extended regions. In contrast, lncRNAs exhibited almost no correlation between length and accessibility in the native condition (P = 0.24; Supplementary Fig. 4i) but a significant positive association after fixation (P = 0.0033; Supplementary Fig. 4j). Because fixation preserves RBP–RNA interactions, this result suggests that length-dependent accessibility in lncRNAs might primarily reflect protein binding rather than intrinsic folding. Together, these findings reveal that accessibility landscapes differ across RNA types and are shaped by both structural and protein-mediated contributions.

### RAD-seq accessibility profile reveals RNA structural changes in regions near modifications

RAD-seq also captured accessibility modulation by RNA chemical modifications. *N*^6^-methyladenosine (m^6^A) sites exhibited increased native accessibility in their flanking regions (P < 2.0 × 10⁻^16^; Supplementary Fig. 5a), consistent with local weakened base pairing^39^. In contrast, regions near pseudouridine (Ψ) modifications, which are generally believed to stabilize RNA secondary structures^40,41^, showed decreased native accessibility (P = 6.4 × 10⁻^6^; Supplementary Fig. 5b). We also observed significantly decreased native accessibility in regions near 5-methylcytidine (m^5^C) sites (P = 1.6 × 10⁻^12^; Supplementary Fig. 5c) this could represent the averaged effects of modifications at NSUN2 and NSUN6 sites^42^, which have different contributions to RNA structure. RAD-seq accessibility can thus be used to detect local structural remodeling caused by the expected biophysical effects of distinct RNA modifications.

### Functional implications of RNA accessibility under cellular stress

Environmental stress rapidly reorganizes RNA metabolism through coordinated changes in transcription, translation, and RNA–protein assembly. Oxidative stress and heat shock both suppress translation but act through distinct pathways that alter RNA structure, RBP engagement, and localization^43–46^. Because RAD-seq simultaneously measures RNA structure and the RBP occupancy of individual adenosines, it offers a unified view of how these stresses remodel the transcriptome. We induced oxidative stress in HepG2 cells with sodium arsenite and heat shock by incubation at 42 °C.

We investigated the global accessibility changes under stress conditions using native RAD-seq as it captures the combined dynamics of RBP binding and RNA structural remodeling *in vivo*. Oxidative stress caused a widespread decrease in native RNA accessibility across both cytosolic and nuclear transcripts (Fig. 3a), consistent with increased RNP assembly and reduced helicase activity during translational inhibition. In contrast, heat shock produced a global increase in native accessibility across both compartments (Fig. 3b), reflecting chaperone-driven remodeling of RNA–protein complexes and changes in RNA structural dynamics. Gini index analyses corroborated these opposing trends and confirmed that transcriptome-wide accessibility distributions shift significantly under stress (Supplementary Fig. 6a-b).

**Figure 3.**
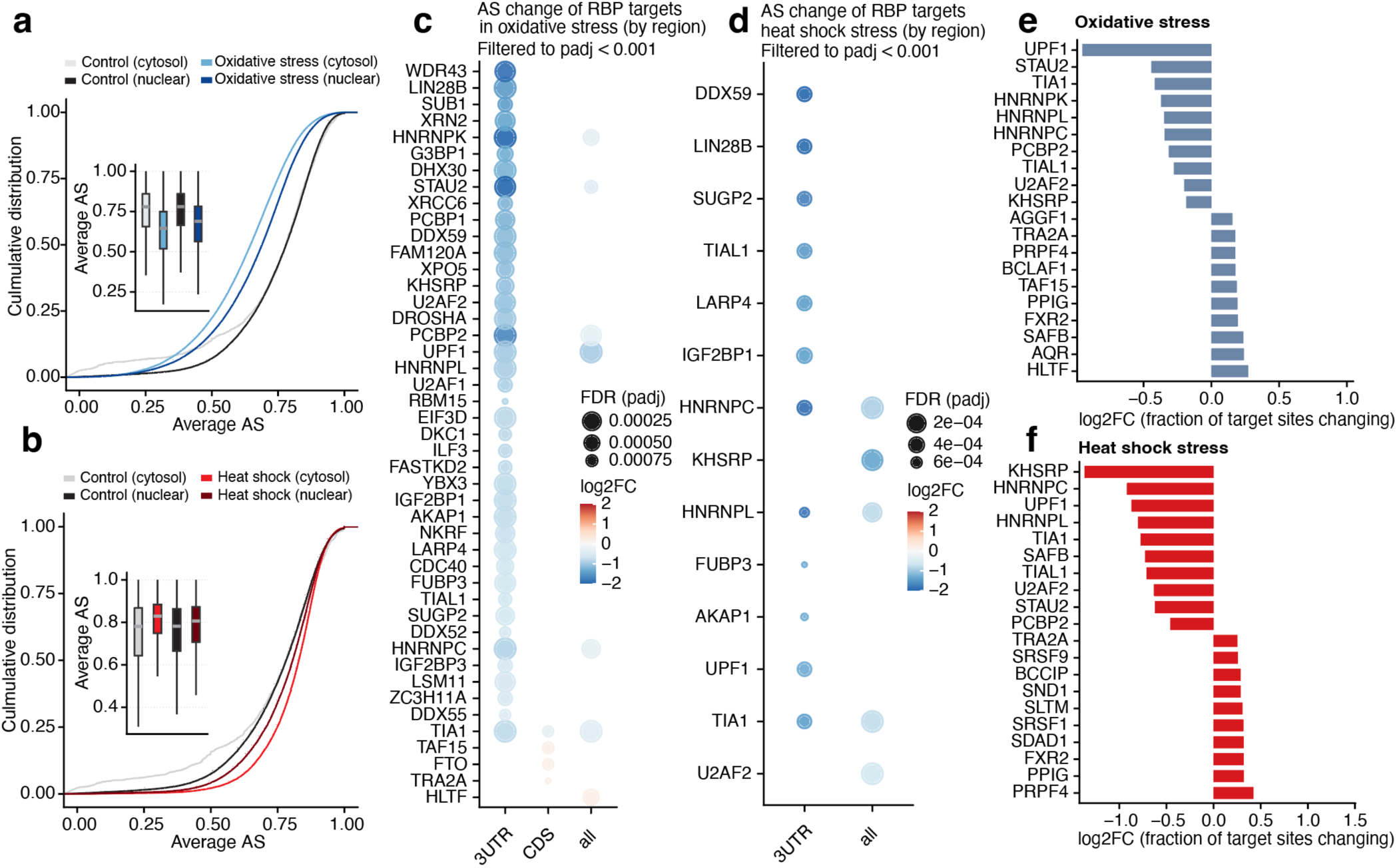
RAD-seq in cellular stress conditions. (a-b) Global RNA accessibility change revealed by RAD-seq in oxidative stress (a) and heat shock stress (b). (c-d) Significantly changed RBP binding targets (p adjusted < 0.001) identified by RAD-seq and ENCODE eCLIP data in HepG2 cells for different regions (3’UTR, CDS, and whole transcript) upon oxidative stress (c) or heat shock stress (d). (e-f) Top 10 RBPs ranked by accessibility change (both positive and negative) of targets in oxidative stress (e) or heat shock stress (f).

To determine how these global signatures relate to specific regulatory factors, we examined native accessibility changes at eCLIP-defined RBP binding sites. Oxidative stress produced significant reductions in accessibility at targets of numerous RBPs, including UPF1, TIA1, HNRNPK, HNRNPL, and IGF2BP proteins (Fig. 3c). These decreases were often strongest in 3′ UTRs, consistent with stabilized RBP occupancy during oxidative stress. RBPs with the largest accessibility reductions at their targets included UPF1, STAU2, and multiple hnRNPs (Fig. 3e), factors involved in mRNA decay, translational repression, and higher-order RNP assembly, which may suggest these RBPs bind to their targets to facilitate RNA degradation and repress translation in response to oxidative stress.

In contrast, heat shock affected a smaller set of RBPs but showed distinct directionality. Although global transcriptome accessibility increased, local accessibility decreased at numerous RBP targets (Fig. 3d), including HNRNPC, KHSRP, IGF2BP1, and DDX59, consistent with sequestration of these mRNAs into stress granules in which transcripts are densely coated by RBPs^47^. The RBPs with the strongest target-associated accessibility increases included splicing-related proteins like SRSF3 and SRSF1 (Fig. 3f), which might imply a decrease in binding and splicing events during heat shock stress. These patterns indicate that heat shock increases accessibility globally through chaperone-driven RNP remodeling yet simultaneously produces marked local accessibility reductions in the subset of transcripts that enter condensed granule-like assemblies.

Together, these results show that RAD-seq resolves stress-dependent remodeling of RNA accessibility at both global and RBP-specific levels. Oxidative stress reduces accessibility transcriptome-wide and strengthens binding of regulatory RBPs, consistent with the formation of compact mRNP assemblies during translational inhibition. Heat shock produces a more complex signature: accessibility increases globally across cytosolic and nuclear transcripts, yet many RBP binding sites show decreased accessibility due to localized condensation of mRNAs into stress granules. By simultaneously reporting global accessibility and local RBP-associated changes from the same experiment, RAD-seq reveals how distinct stress pathways differentially reshape RBP engagement across the transcriptome.

### RBP-specific accessibility footprints revealed by RAD-seq

The transcriptome-wide analyses above established that RAD-seq captures meaningful variation in RNA accessibility driven by structure, modification, and cellular state. Our observation of stress-induced accessibility shifts at known binding sites for specific RBPs further indicated that RAD-seq profiles may encode individual protein-binding footprints that reflect intrinsic properties of individual RBPs.

To test this possibility, we first generated length-normalized meta-footprints centered on ENCODE eCLIP peaks by the ENCODE project in HepG2 cells^48^. Although the specific repertoire of target transcripts differs between cell types, the stereochemical features of RBP-RNA interactions and the structural footprints imposed by an RBP’s physical dimensions are intrinsic to the protein itself^49^. Thus, high-confidence HepG2 binding sites provide suitable proxies for aggregating and resolving the characteristic structural geometries of these RBPs in the HeLa context (Fig. 4a). Across RBPs, fixed accessibility from HeLa cells was reduced at protein binding regions relative to the background transcriptome (Fig. 4b), and consistently lowest at peak centers, with increased fixed accessibility in the surrounding regions (Fig. 4c), matching the expected signature of protein protection at binding sites. icSHAPE profiles displayed similar, though weaker, trends (Fig. 4d-f), suggesting that RAD-seq robustly detects features of RNA–RBP interactions that are only partially captured by conventional chemical probing (including RBP occlusion and the structural contexts that favor binding).

**Figure 4.**
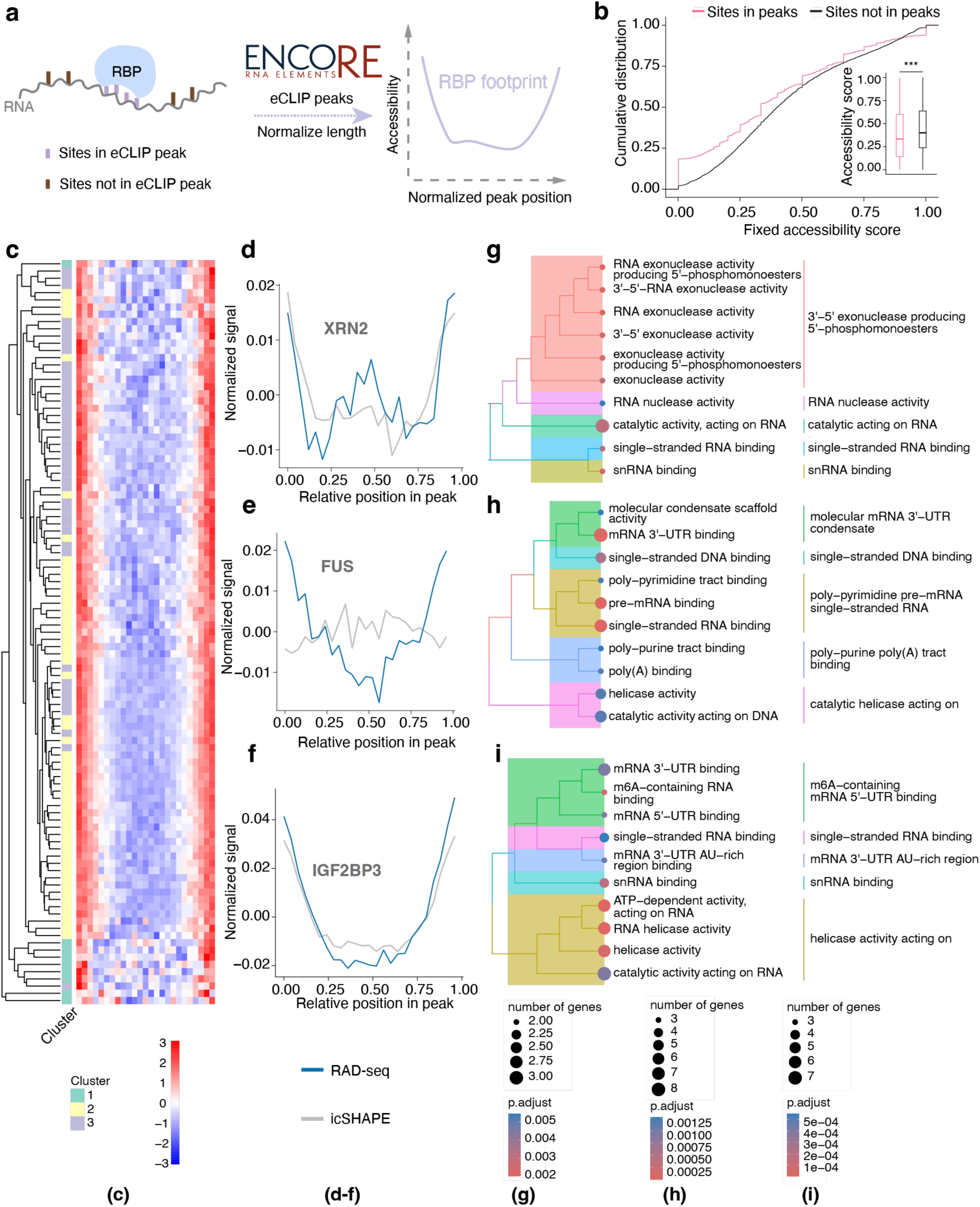
Identification of RBP clusters using RNA accessibility profile. (a) Schematic of RBP footprinting. For each of 103 RBPs in HepG2 (ENCODE eCLIP), peaks were scaled to a common length, and the mean fixed RAD-seq accessibility from HeLa cells across the scaled peaks was plotted to reveal a characteristic footprint. (b) Fixed accessibility at sites within eCLIP peaks (pink) versus non-peak sites (black); groups compared with a two-sided Wilcoxon rank-sum test (P < 0.01). (c) Heatmap of normalized accessibility profiles for 103 HepG2 RBPs; each row is z-scored and clustered by K-means clustering (k = 3). (d–f) Average fixed RAD-seq signal (blue) and icSHAPE signal (gray), mean-centered within normalized peaks, for clusters 1 (d), 2 (e), and 3 (f). (g–i) Gene Ontology Molecular Function tree plots for RBP clusters 1 (g), 2 (h), and 3 (i).

Next, we sought to examine whether distinct RBPs that exhibit characteristic accessibility signatures could be identified by their meta-footprints. Hierarchical clustering of RBP-specific meta-footprints revealed three reproducible classes (Fig. 4c). Class 1 showed heterogeneous accessibility at the peak center with variable shoulders; Class 2 displayed deep, symmetric valleys; and Class 3 exhibited broad, shallow depressions extending into the flanks (Fig. 4d–f). Functional enrichment analysis indicated clear biological coherence. Class 1 RBPs were enriched for exonuclease and 3′/5′ UTR-associated factors (Fig. 4g), consistent with the transient or distributive interactions suggested by their RAD-seq signatures. Class 2 contained canonical mRNA 3′ UTR binders and single-stranded RNA-binding proteins (Fig. 4h), reflecting stable engagement with accessible regions. Class 3 was enriched for catalytic RBPs, including RNA helicases and m^6^A-recognition proteins (Fig. 4i), suggesting extended remodeling or multivalent interactions.

Together, these analyses demonstrate that RAD-seq accessibility encodes rich, RBP-specific binding information, revealing stereotyped footprint shapes that align with eCLIP-defined binding sites and correspond to mechanistically distinct classes of RNA–protein interaction. RAD-seq thus converts accessibility profiles into a high-resolution atlas of in vivo RBP occupancy, bridging global accessibility landscapes with local molecular determinants.

### Motif-guided inference of RBP binding changes from accessibility profiles

Conventional CLIP-seq requires one experiment per RBP and often suffers from variable resolution and signal-to-noise. Because RAD-seq directly reports RNA accessibility, which is reduced by protein occlusion, we asked whether accessibility data, when combined with known motif preferences, could be used to infer RBP binding changes without additional immunoprecipitation experiments.

For RBPs with available RNA Bind-n-Seq (RBNS) motifs^50^, we defined a protein-binding score (PBscore) at each motif instance as the difference between native and fixed accessibility (PBscore = Native AS – Fixed AS), where larger positive values indicate stronger protein protection. If an RBP occupies its preferred motifs *in vivo*, these sites should exhibit lower fixed accessibility and thus higher protein-binding scores.

As a proof of concept, we evaluated whether RAD-seq captures sequence preferences of specific RBPs by comparing its accessibility ranks to RBFOX2 binding enrichment measured by RBNS across all 256 adenosine-centered 5-mer motifs. We observed a strong negative correlation between motif accessibility in the fixed condition and RBNS enrichment (Supplementary Fig. 7a–d), showing that RBFOX2-preferred motifs are less accessible *in vivo*, consistent with direct protein binding.

We next applied this approach to quantify condition-dependent binding changes following knockdown of EIF4A3, which disrupts exon junction complex (EJC) assembly. The EJC normally restricts access to nearby RNA sites; therefore, EIF4A3 depletion is expected to increase local accessibility and reduce PBscores at these positions. Conversely, proteins whose binding is unmasked by EJC loss should show positive change in PBscore. For each RBP, we ranked motif instances by the change in PBscore (ΔPBscore) between wild-type and knockdown cells and performed gene set enrichment analysis (GSEA) ^51^ against the corresponding RBNS motif sets (Fig. 5a). This analysis identified TRA2A as the top-enriched RBP with elevated ΔPBscore concentrated in EJC-associated regions (Fig. 5b–e; Supplementary Figs. 8,9). Notably, TRA2A has been shown to interact with METTL3^52^, a catalytic component of the m^6^A methyltransferase complex. These observations suggest a previously unrecognized potential pathway in which loss of EIF4A3 may expose occluded sites near EJCs, thereby enabling TRA2A recruitment of METTL3 and facilitating local m^6^A deposition (Fig. 5f).

**Figure 5.**
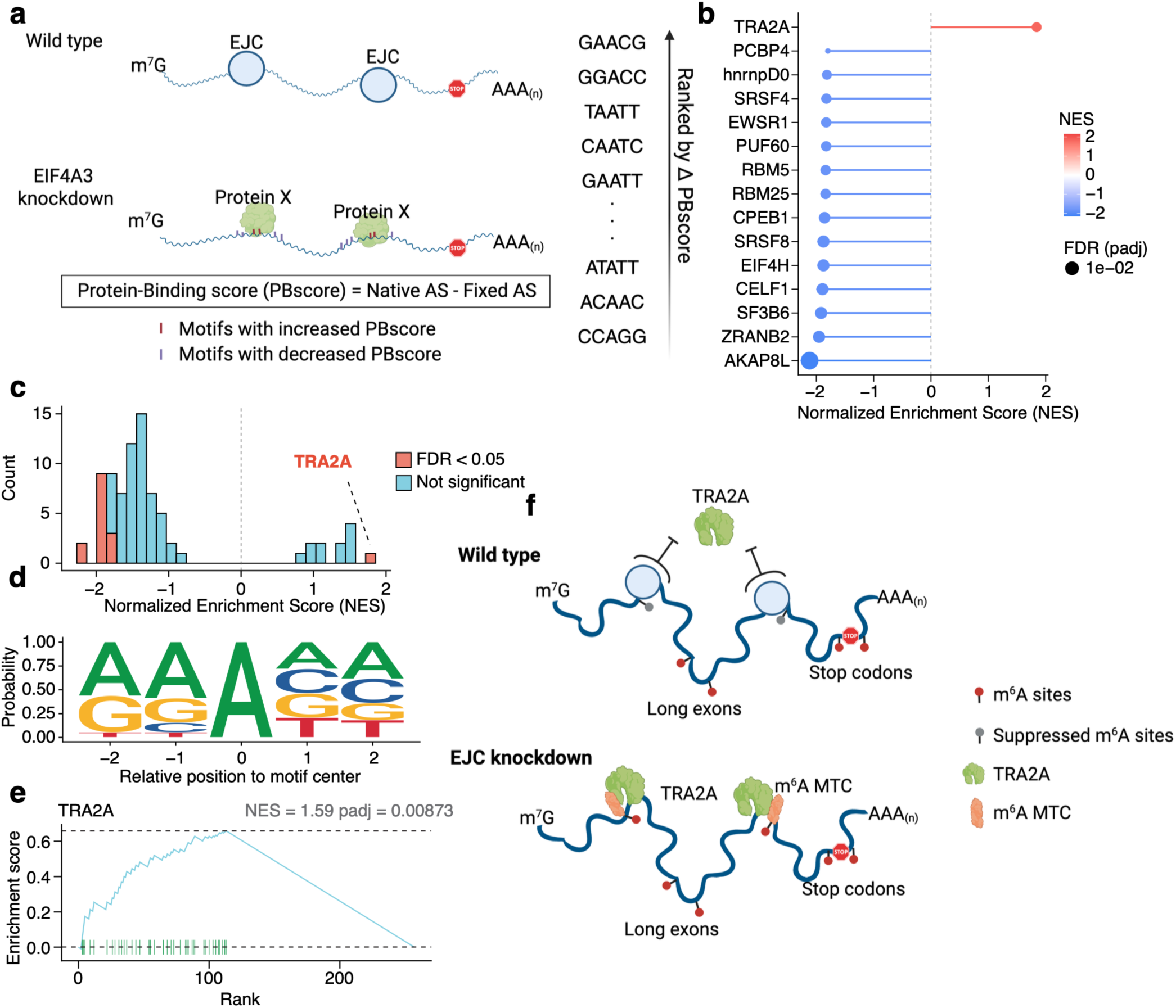
Inferring RBP binding from RBNS motifs and RAD-seq accessibility changes. (a) Schematic of the inference framework. RBNS-derived preferred k-mers for each RBP were used as candidate motifs; motif occurrences overlapping EIF4A3 eCLIP peaks from HeLa cells were evaluated by the RAD-seq protein-binding score from HeLa cells and ranked by the change between EIF4A3 knockdown and wild type (ΔPBscore = PBscore_KD − PBscore_WT). Created in BioRender. Li, Y. (2025) https://BioRender.com/9vv6474. (b) RBPs with the twenty highest normalized enrichment scores (NES) from a GSEA testing whether each RBP’s RBNS-preferred k-mers are enriched at the top of the ΔPBscore-based motif ranking. (c) NES distribution across all RBPs; TRA2A is the only RBP with significant positive enrichment (NES > 0, FDR < 0.05; Benjamini–Hochberg adjustment). (d) Aggregate motif-centered RAD-seq profile for TRA2A using its top sixty RBNS-preferred k-mers aligned at the central adenosine. (e) GSEA running-enrichment plot for TRA2A showing the locations of preferred k-mers in the list of motifs ranked by PBscore, along with the NES and the Benjamini–Hochberg adjusted P value. (f) Proposed model. In wild type, the exon junction complex anchored by EIF4A3 excludes TRA2A and the METTL3–METTL14 methyltransferase complex from exon junction regions; after EIF4A3 knockdown, TRA2A binds near the junction and recruits the methyltransferase complex, increasing m^6^A deposition at these sites. Created in BioRender. Li, Y. (2025) https://BioRender.com/jr9d511.

Together, these results demonstrate that RAD-seq, combined with simple motif priors, can quantitatively infer condition-dependent RBP binding dynamics from a single accessibility dataset. This approach recapitulates information typically requiring numerous CLIP-seq experiments, establishing RAD-seq as a scalable platform for transcriptome-wide discovery of dynamic protein–RNA interactions.

### Model-based inference of RBP occupancy from RAD-seq accessibility

The motif-guided perturbation analysis above established that accessibility changes encode information about protein binding dynamics. We next asked whether RAD-seq accessibility alone, integrated with sequence context, could predict RBP occupancy transcriptome-wide without per-RBP experiments.

To this end, we developed an unsupervised statistical framework that combines RAD-seq accessibility with known RBP binding motifs to infer *in vivo* occupancy from a single dataset (Fig. 6a). Conceptually similar to CENTIPEDE^53^ for transcription factors, the model integrates a motif-informed prior with an accessibility-based footprint likelihood, jointly estimating whether each motif instance is bound or unbound. For every RBP, we scan the transcriptome using position weight matrices (PWMs) from the CISBP-RNA database^54^ and evaluate motif occurrences within their local accessibility context. The model treats the RAD-seq signal around each motif as a mixture of two latent states—bound and unbound— capturing the expected depletion of accessibility at bound sites.

**Figure 6.**
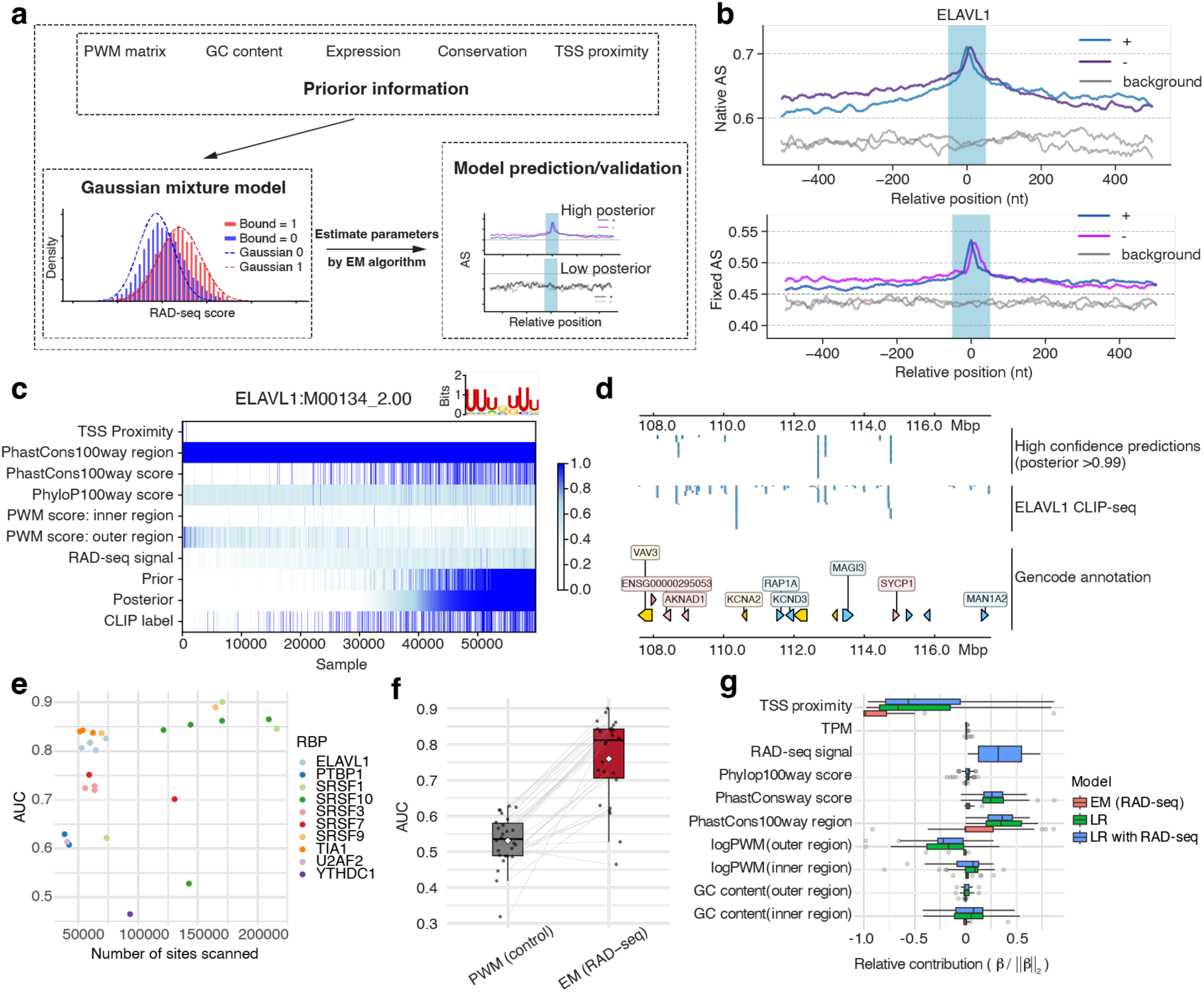
Unsupervised statistical model for RBP binding-site inference using RBP motifs in HeLa cells. (a) Overview of the model integrating known RBP motifs with RAD-seq accessibility to infer binding probabilities. The prior combines CISBP-RNA PWM scores computed in inner (motif-centered) and outer flanking windows, GC content, RNA-seq expression, distance to the TSS, and conservation (PhastCons100way and PhyloP100way). A RAD-seq–derived accessibility statistic (RAD-seq data from HeLa cell, see Methods) is modeled with a two-component Gaussian mixture representing bound and unbound states, and parameters as well as posterior binding probabilities are estimated by expectation maximization (EM). (b) Accessibility profiles centered on ELAVL1 binding sites showing higher signal within 50 bp of the site relative to distal regions (>50 bp); gray traces indicate matched genomic background. (c) Heatmap of features and outputs for ELAVL1 using CISBP-RNA motif M00134_2, including TSS proximity, PhastCons100way, PhyloP100way, PWM score (inner window), PWM score (outer window), RAD-seq statistic, prior, posterior, and CLIP-seq label. (d) IGV example illustrating concordance between high-confidence predictions (posterior > 0.99) and ELAVL1 CLIP-seq peaks. (e) Area under the ROC curve (AUC) as a function of the number of scanned sites (n = 9 RBPs from POSTAR HeLa cell CLIP-seq; 30 motifs). (f) Performance comparison of the EM model that includes RAD-seq features versus a PWM-only baseline. (g) Normalized coefficient contributions 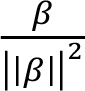 from the EM model with RAD-seq, logistic regression (LR) without RAD-seq, and LR with RAD-seq.

Priors incorporate PWM score, GC content, gene expression level, sequence conservation, and distance to the transcription start site, while the likelihood is derived directly from RAD-seq, summarizing accessibility in a 50-nt central window and 100-nt flanking regions around each motif. A two-component Gaussian mixture model, fitted by expectation–maximization, outputs posterior probabilities of RBP binding for every motif instance (Fig. 6b).

Using ELAVL1 with the M00134_2 motif as an example, the model recovered binding sites that strongly overlapped known CLIP-seq peaks from POSTAR^55^ (Fig. 6c,d). Across nine RBPs and thirty motifs, the framework achieved a median AUC > 0.8, outperforming PWM-only baselines (Fig. 6e,f). Feature attribution analysis revealed that sequence conservation and TSS proximity contributed most to predictive power, with PWM score and expression providing additional signal (Fig. 6g).

Together, these results demonstrate that RAD-seq accessibility profiles, combined with sequence-based priors, are sufficient to deconvolute RBP occupancy at scale. This model transforms a single RAD-seq experiment into transcriptome-wide binding maps for dozens of RBPs, providing an efficient alternative to large-scale CLIP-seq profiling and other targeted assays for large-scale profiling of RNA–protein interaction landscapes.

## Discussion

We developed RAD-seq, an enzymatic assay that measures RNA accessibility *in vivo* through TadA8.20-mediated adenosine deamination in a simple, scalable, and cost-effective workflow. RAD-seq reports a unified signal that accurately reflects both structural openness and protein occupancy, and it does so at single-nucleotide resolution from a simple workflow. Across multiple analyses, accessibility scores aligned with known structural features, captured expected effects of RNA modifications, and revealed reproducible RBP footprints, establishing RAD-seq as a quantitative, interpretable, and generalizable framework for transcriptome-wide accessibility mapping, analogous to how ATAC-seq or DNA enzymatic methylation assays define chromatin openness.

Beyond structural information, RAD-seq can report on regulatory states that are difficult to resolve with existing RNA assays. Although chemical probe-based measurements (SHAPE, DMS, and LASER) are precise at the nucleotide resolution, which is essential to map structures of target RNA, RAD-seq measures the RBP protection of RNA at the scale of protein-binding level instead of solvent accessibility. DNA 5mC methylation and 6mA methylation approach have been successfully developed and employed to map DNA accessibility in the past^32–36^. RAD-seq is the equivalent for RNA but relying on base editing to facilitate more straightforward readout.

Because accessibility reflects both folding and RBP engagement, RAD-seq sensitively detects transcriptome-wide remodeling of RNA organization during cellular perturbations. Under oxidative and heat-shock stress, the assay distinguished global shifts in RNA openness from localized reductions at RBP-bound regions, suggesting how stress granule condensation and RNP remodeling reshape the transcriptome. Together, these features illustrate RAD-seq’s potential to connect biophysical RNA states with regulatory outcomes, helping to map both RNA structure and RBP protection.

A key advantage of RAD-seq is the potential to infer RBP occupancy at scale from a single dataset. By integrating motif priors with an unsupervised statistical model that combines native and fixed accessibility, RAD-seq can recover motif-resolved footprints and generate transcriptome-wide binding landscapes without per-RBP profiling. This framework enables rapid and cost-effective surveys of condition-dependent RBP-associated accessibility changes and can help prioritize candidates for follow-up with eCLIP or orthogonal assays.

The interpretive potential of RAD-seq likely extends beyond the contexts explored here. Because accessibility decreases wherever RNA surfaces become occluded, RAD-seq can potentially identify endogenous RNA–RNA interactions by detecting local protection patterns arising from long-range intramolecular folding or intermolecular pairing. Moreover, applied to chromatin-associated RNAs, RAD-seq could also help clarify how transcriptional states, co-transcriptional RNP assembly, and chromatin architecture jointly shape RNA organization at gene loci. These possibilities position RAD-seq not merely as an RNA structure assay but also as a more general platform for probing higher-order RNA interactions and chromatin-tethered regulation.

## Limitations

Like all enzymatic accessibility assays, RAD-seq has technical limitations. TadA8.20 edits only adenosines and shows reduced activity at m^6^A, making accessibility estimates sensitive to sequence context, RNA abundance, and sequencing depth. Like ATAC-seq and other enzymatic methods, TadA8.20 does have sequence preferences that in principle can be calibrated. Formaldehyde fixation may also introduce accessibility biases, and RBP-occupancy inference depends on motif libraries that remain incomplete. Improvements in background modeling, enzyme engineering, and complementary chemistries, together with expanded motif resources and additional benchmarking, will further increase accuracy. As these improvements mature, RAD-seq will become an increasingly powerful and scalable platform for quantitative, transcriptome-wide mapping of RNA structure and RBP engagement *in vivo*.

## Methods

### Overexpression and purification of recombinant TadA8.20 protein from the EM model

TadA8.20 plasmids were obtained from Addgene (Plasmid #194702). BL21 Rosetta 2 (DE3) competent cells were transformed with the recombinant plasmids and grown on Luria broth (LB) agar plates supplemented with 50 μg/mL kanamycin and 25 μg/mL chloramphenicol. Successfully transformed bacteria were always cultured in the presence of 50 μg/mL kanamycin and 25 μg/mL chloramphenicol unless otherwise noted. Single colonies were inoculated into fresh LB medium and grown in an incubator shaker (37°C, 220 rpm) for 12–18 h. A 10 mL saturated start culture was used to inoculate 1 L fresh medium. Bacteria were grown at 37°C until OD_600_ reached 0.5. The culture was cooled down immediately to 4°C and induced with 0.1 mM isopropyl β-d-1-thiogalactopyranoside (IPTG). Bacteria were cultured at 16°C for an additional 20 h before pelleting by centrifugation at 4,000 g.

Bacterial pellets were lysed by sonication in buffer A (50 mM Tris, 500 mM NaCl, 10 mM β-mercaptoethanol, and 10% (v/v) glycerol; pH 7.5). Lysed bacteria were clarified by centrifugation at 4°C, 23,000 g. The supernatant was loaded onto a Ni-NTA Superflow Cartridge (Qiagen, 30761), washed with 30 mL of buffer A supplemented with 50 mM imidazole, and eluted with a gradient of imidazole from 50 mM to 500 mM in buffer A.

The eluted protein was incubated with TEV protease and dialyzed in buffer A at 4°C overnight. The protein mixture was reloaded onto a Ni-NTA Superflow Cartridge, washed with buffer B (50 mM Tris, 1 M NaCl, 10 mM β-mercaptoethanol, and 10% (v/v) glycerol; pH 8.0), and eluted by buffer B supplemented with 50 mM imidazole. Finally, MBP-free TadA8.20 was purified by size-exclusion chromatography (Enrich^™^ SEC 650 10 × 300 mm Column, Bio-Rad, 7801650) and concentrated to approximately 4 mg/mL. The column was balanced and eluted with buffer C (50 mM Tris, 200 mM NaCl, 10 mM β-mercaptoethanol, and 10% (v/v) glycerol; pH 7.5).

### Cell culture

HeLa and HepG2 cell lines were grown in Dulbecco’s modified Eagle’s medium (DMEM; Gibco, catalog no. 11965092) supplemented with 10% fetal bovine serum (FBS; Gibco) and 1% 100× penicillin–streptomycin (Gibco).

### Library preparation for RAD-seq

After deaminase labeling, ribosomal RNA (rRNA) was depleted by RiboMinus rRNA depletion kits (Thermo) following the manufacturer’s protocol. For cDNA library construction, RNA samples were mixed in a 10 µL reverse transcription reaction. For cDNA library construction, RNA samples were mixed in a 10 µL reverse transcription reaction. Each reaction contained 7 µL RNA and nuclease-free water, 1 µL 100 µM s5-primer mix (prepared as 33 µM s5-dT30VN, 5′-ACACTCTTTCCCTACACGACGCTCTTCCGATCTAGACTCNNNNNNTTTTTTTTTTTTTTTTTTTTTTTT TTTTTTTVN-3′; 33 µM s5-N6, 5′-ACACGACGCTCTTCCGATCTNNNNNN-3′; and 33 µM s5-V6, 5′-ACACGACGCTCTTCCGATCTVVVVVV-3′), 2 µL 50 percent PEG 8000, and 1 µL 10 mM dNTPs. The mixture was incubated at 65 °C for 5 min and cooled on ice. After this, 1 µL s7-TSO (5′-/5SpC3/CAGACGTGTGCTCTTCCGATCTNNNNNNrGrGrG-3′), 4 µL 5× Maxima H Minus reverse transcription buffer (Thermo, EP0752), 2 µL 10 mM GTP, 1 µL RNaseOUT (Thermo, 10777019), and 1 µL Maxima H Minus reverse transcriptase (Thermo, EP0752) were added. Reverse transcription was carried out at 42 °C for 90 min, followed by ten cycles of 50 °C for 2 min and 42 °C for 2 min, and a final incubation at 85 °C for 5 min to inactivate the enzyme. cDNA products were purified using 0.8× AMPure XP beads according to the manufacturer’s instructions, eluted in nuclease-free water, and used for PCR amplification. Libraries were amplified with 12 PCR cycles, followed by a second 0.8× AMPure XP purification. Size selection of fragments larger than 300 base pairs was performed using low-melting agarose, and DNA was recovered using three volumes of Buffer QG relative to the gel mass.

### Data preprocessing of RAD-seq

Adapters were removed from raw data by Cutadapt^57^. Reads were first mapped to human rRNA sequences using HISAT-3N^58^ (--base-change A,G –no-spliced-alignment –no-softclip). The remaining non-rRNA reads were mapped to the human genome (hg38) and the GENCODE (GRCh38.p14) transcript annotation^59^ (--time --base-change A,G --no-unal). The mapped reads were deduplicated by the subcommand dedup from UMI-tools^60^. Sites were filtered with depth >=3 and the accessibility scores were calculated by the converted ratio from A to G. The Gini index^61^ was calculated in each transcript or UTR/CDS by annotation from GENCODE gtf file^59^.

### Induction of stress conditions

Heat shock stress was induced by replacing the cell culture media with fresh media and incubating the cells at 42 °C for the intended time (1 h or 2 h) before harvesting. Oxidative stress was induced by replacing the cell culture media with fresh media supplemented with 0.5 mM sodium arsenite and incubating the cells at 37 °C for 1 h before harvesting. Recovery was performed by replacing the cell culture media with fresh media and incubating the stress-induced cells at 37 °C for 1h before harvesting.

## Acknowledgements

The authors thank Drs. Weixin Tang, Yuan Wu, and Dr. Yulan Xiao for providing the TadA8.20 used in our experiments. We thank P. Faber of the University of Chicago Genomics Facility for sequencing support. The authors thank Natalia Gonzales and Fred Naumann for their comments and edits, which significantly improved the manuscript. C.H. is an Investigator of the Howard Hughes Medical Institute.

## Data availability

All high-throughput sequencing data generated in this study have been deposited in the NCBI Gene Expression Omnibus (GEO) under accession number GSE311901.

## Reference

1. Boyle, A. P. et al. High-Resolution Mapping and Characterization of Open Chromatin across the Genome. Cell 132, 311–322 (2008).

2. Buenrostro, J. D., Giresi, P. G., Zaba, L. C., Chang, H. Y. & Greenleaf, W. J. Transposition of native chromatin for fast and sensitive epigenomic profiling of open chromatin, DNA-binding proteins and nucleosome position. Nat. Methods 10, 1213–1218 (2013).

3. Weng, X. et al. Keth-seq for transcriptome-wide RNA structure mapping. Nat. Chem. Biol. 16, 489–492 (2020).

4. Rouskin, S., Zubradt, M., Washietl, S., Kellis, M. & Weissman, J. S. Genome-wide probing of RNA structure reveals active unfolding of mRNA structures in vivo. Nature 505, 701–705 (2014).

5. Ding, Y. et al. In vivo genome-wide profiling of RNA secondary structure reveals novel regulatory features. Nature 505, 696–700 (2014).

6. Lucks, J. B. et al. Multiplexed RNA structure characterization with selective 2′-hydroxyl acylation analyzed by primer extension sequencing (SHAPE-Seq). Proc. Natl. Acad. Sci. 108, 11063–11068 (2011).

7. Spitale, R. C. et al. Structural imprints in vivo decode RNA regulatory mechanisms. Nature 519, 486–490 (2015).

8. Zubradt, M. et al. DMS-MaPseq for genome-wide or targeted RNA structure probing in vivo. Nat. Methods 14, 75–82 (2017).

9. Siegfried, N. A., Busan, S., Rice, G. M., Nelson, J. A. E. & Weeks, K. M. RNA motif discovery by SHAPE and mutational profiling (SHAPE-MaP). Nat. Methods 11, 959–965 (2014).

10. Luo, Q.-J. et al. RNA structure probing reveals the structural basis of Dicer binding and cleavage. Nat. Commun. 12, 3397 (2021).

11. Zinshteyn, B. et al. Assaying RNA structure with LASER-Seq. Nucleic Acids Res. 47, 43–55 (2019).

12. Chan, D. et al. Diverse functional elements in RNA predicted transcriptome-wide by orthogonal RNA structure probing. Nucleic Acids Res. 49, 11868–11882 (2021).

13. Wan, Y., Qu, K., Ouyang, Z. & Chang, H. Y. Genome-wide mapping of RNA structure using nuclease digestion and high-throughput sequencing. Nat. Protoc. 8, 849–869 (2013).

14. Kertesz, M. et al. Genome-wide measurement of RNA secondary structure in yeast. Nature 467, 103–107 (2010).

15. Underwood, J. G. et al. FragSeq: transcriptome-wide RNA structure probing using high-throughput sequencing. Nat. Methods 7, 995–1001 (2010).

16. Silverman, I. M. & Gregory, B. D. Transcriptome-wide ribonuclease-mediated protein footprinting to identify RNA–protein interaction sites. Methods 72, 76–85 (2015).

17. Silverman, I. M. et al. RNase-mediated protein footprint sequencing reveals protein-binding sites throughout the human transcriptome. Genome Biol. 15, R3 (2014).

18. Ule, J., Jensen, K., Mele, A. & Darnell, R. B. CLIP: A method for identifying protein–RNA interaction sites in living cells. Methods 37, 376–386 (2005).

19. Johnson, D. S., Mortazavi, A., Myers, R. M. & Wold, B. Genome-Wide Mapping of in Vivo Protein-DNA Interactions. Science 316, 1497–1502 (2007).

20. Huppertz, I. et al. iCLIP: Protein–RNA interactions at nucleotide resolution. Methods San Diego Calif 65, 274–287 (2014).

21. Van Nostrand, E. L. et al. Robust transcriptome-wide discovery of RNA-binding protein binding sites with enhanced CLIP (eCLIP). Nat. Methods 13, 508–514 (2016).

22. Weidmann, C. A., Mustoe, A. M., Jariwala, P. B., Calabrese, J. M. & Weeks, K. M. Analysis of RNA–protein networks with RNP-MaP defines functional hubs on RNA. Nat. Biotechnol. 39, 347–356 (2021).

23. Meyer, K. D. DART-seq: an antibody-free method for global m6A detection. Nat. Methods 16, 1275–1280 (2019).

24. Xu, W., Rahman, R. & Rosbash, M. Mechanistic implications of enhanced editing by a HyperTRIBE RNA-binding protein.

25. Simultaneous profiling of the RNA targets of two RNA-binding proteins using TRIBE-STAMP. in Methods in Enzymology vol. 705 127–157 (Elsevier, 2024).

26. McMahon, A. C. et al. TRIBE: Hijacking an RNA-Editing Enzyme to Identify Cell-Specific Targets of RNA-Binding Proteins. Cell 165, 742–753 (2016).

27. Brannan, K. W. et al. Robust single-cell discovery of RNA targets of RNA-binding proteins and ribosomes. Nat. Methods 18, 507–519 (2021).

28. Flamand, M. N., Ke, K., Tamming, R. & Meyer, K. D. Single-molecule identification of the target RNAs of different RNA binding proteins simultaneously in cells. Genes Dev. genesdev;gad.349983.122v1 (2022) doi:10.1101/gad.349983.122.

29. Xiao, Y.-L. et al. Transcriptome-wide profiling and quantification of N6-methyladenosine by enzyme-assisted adenosine deamination. Nat. Biotechnol. 41, 993–1003 (2023).

30. Gaudelli, N. M., et al. Directed evolution of adenine base editors with increased activity and therapeutic application. Nat. Biotechnol. 38, 892–900 (2020).

31. Gaudelli, N. M. et al. Programmable base editing of A•T to G•C in genomic DNA without DNA cleavage. Nature 551, 464–471 (2017).

32. Krebs, A. R. et al. Genome-wide Single-Molecule Footprinting Reveals High RNA Polymerase II Turnover at Paused Promoters. Mol. Cell 67, 411–422.e4 (2017).

33. Guo, H. et al. DNA methylation and chromatin accessibility profiling of mouse and human fetal germ cells. Cell Res. 27, 165–183 (2017).

34. Kelly, T. K. et al. Genome-wide mapping of nucleosome positioning and DNA methylation within individual DNA molecules. Genome Res. 22, 2497–2506 (2012).

35. Shipony, Z. et al. Long-range single-molecule mapping of chromatin accessibility in eukaryotes. Nat. Methods 17, 319–327 (2020).

36. Stergachis, A. B., Debo, B. M., Haugen, E., Churchman, L. S. & Stamatoyannopoulos, J. A. Single-molecule regulatory architectures captured by chromatin fiber sequencing. Science 368, 1449–1454 (2020).

37. Sweeney, B. A. et al. R2DT is a framework for predicting and visualising RNA secondary structure using templates. Nat. Commun. 12, 3494 (2021).

38. Andrews, R. J., Baber, L. & Moss, W. N. RNAStructuromeDB: A genome-wide database for RNA structural inference. Sci. Rep. 7, 17269 (2017).

39. Liu, N. et al. N6-methyladenosine-dependent RNA structural switches regulate RNA-protein interactions. Nature 518, 560–564 (2015).

40. Schwartz, S. et al. Transcriptome-wide Mapping Reveals Widespread Dynamic-Regulated Pseudouridylation of ncRNA and mRNA. Cell 159, 148–162 (2014).

41. Carlile, T. M. et al. Pseudouridine profiling reveals regulated mRNA pseudouridylation in yeast and human cells. Nature 515, 143–146 (2014).

42. Liu, J. et al. Developmental mRNA m5C landscape and regulatory innovations of massive m5C modification of maternal mRNAs in animals. Nat. Commun. 13, 2484 (2022).

43. Kharel, P. et al. Stress promotes RNA G-quadruplex folding in human cells. Nat. Commun. 14, 205 (2023).

44. Bresson, S. et al. Stress-Induced Translation Inhibition through Rapid Displacement of Scanning Initiation Factors. Mol. Cell 80, 470–484.e8 (2020).

45. Spriggs, K. A., Bushell, M. & Willis, A. E. Translational Regulation of Gene Expression during Conditions of Cell Stress. Mol. Cell 40, 228–237 (2010).

46. Ivanov, P., Kedersha, N. & Anderson, P. Stress Granules and Processing Bodies in Translational Control. Cold Spring Harb. Perspect. Biol. 11, a032813 (2019).

47. Khong, A. et al. The stress granule transcriptome reveals principles of mRNA accumulation in stress granules. Mol. Cell 68, 808–820.e5 (2017).

48. Van Nostrand, E. L. et al. A large-scale binding and functional map of human RNA-binding proteins. Nature 583, 711–719 (2020).

49. Dominguez, D., et al. Sequence, Structure, and Context Preferences of Human RNA Binding Proteins. Mol. Cell 70, 854–867.e9 (2018).

50. Lambert, N. et al. RNA Bind-n-Seq: Quantitative Assessment of the Sequence and Structural Binding Specificity of RNA Binding Proteins. Mol. Cell 54, 887–900 (2014).

51. Subramanian, A. et al. Gene set enrichment analysis: A knowledge-based approach for interpreting genome-wide expression profiles. Proc. Natl. Acad. Sci. 102, 15545–15550 (2005).

52. An, S. et al. Integrative network analysis identifies cell-specific trans regulators of m6A. Nucleic Acids Res. 48, 1715–1729 (2020).

53. Pique-Regi, R. et al. Accurate inference of transcription factor binding from DNA sequence and chromatin accessibility data. Genome Res. 21, 447–455 (2011).

54. Rosado-Tristani, D. A. et al. CisBP-RNA: a web resource for eukaryotic RNA-binding proteins and their motifs. Nucleic Acids Res. gkaf1081 (2025) doi:10.1093/nar/gkaf1081.

55. Zhao, W. et al. POSTAR3: an updated platform for exploring post-transcriptional regulation coordinated by RNA-binding proteins. Nucleic Acids Res. 50, D287–D294 (2022).

56. Klemm, S. L., Shipony, Z. & Greenleaf, W. J. Chromatin accessibility and the regulatory epigenome. Nat. Rev. Genet. 20, 207–220 (2019).

57. Martin, M. Cutadapt removes adapter sequences from high-throughput sequencing reads. EMBnet.journal 17, 10–12 (2011).

58. Zhang, Y., Park, C., Bennett, C., Thornton, M. & Kim, D. Rapid and accurate alignment of nucleotide conversion sequencing reads with HISAT-3N. Genome Res. 31, 1290–1295 (2021).

59. Mudge, J. M., et al. GENCODE 2025: reference gene annotation for human and mouse. Nucleic Acids Res. 53, D966–D975 (2025).

60. Smith, T., Heger, A. & Sudbery, I. UMI-tools: modeling sequencing errors in Unique Molecular Identifiers to improve quantification accuracy. Genome Res. 27, 491–499 (2017).

61. Jiang, L., Chen, H., Pinello, L. & Yuan, G.-C. GiniClust: detecting rare cell types from single-cell gene expression data with Gini index. Genome Biol. 17, 144 (2016).

62. Ray, D. et al. A compendium of RNA-binding motifs for decoding gene regulation. Nature 499, 172–177 (2013).

